# Environmental enrichment accelerates cortical memory consolidation

**DOI:** 10.1101/2025.07.28.667332

**Authors:** Ingrid M. Esteves, HaoRan Chang, Adam R. Neumann, Bruce L. McNaughton

## Abstract

Environmental enrichment is an established strategy to enhance learning and to build resilience against neurodegeneration. In humans, this is known as ‘cognitive reserve’. Though the beneficial effects of exposure to a complex environment in animal models have been well-documented by behavioural, immunohistological and morphological observations, its impact on the functional properties of neuronal populations remains poorly understood. This study aimed to compare the functional encoding and offline memory retrieval dynamics of cortical neurons in enriched and control mice performing a virtual spatial foraging task. Thy1-GCaMP6s mice aged 21 days were enriched for 9 weeks by running a complex obstacle course, during which they were gradually exposed to many different types of obstacles requiring climbing, jumping, and/or balancing elements. Control animals were exercise-matched by running a similar track containing only repeating ramps. At the age of 3 months, two-photon calcium imaging was conducted on populations of neurons from the secondary motor cortex in both groups before, during, and after repeated locomotion through a virtual environment with salient visual-tactile cues. We observed an increase in memory reactivation in the enriched group during the first day of exposure. With training, enriched animals exhibited a stronger anticipatory reduction in running speed near the reward location. Moreover, cortical neuron activity representing locations on the track became substantially more stable and precise over days in enriched but not control animals. Altogether, these results indicate that prior environmental enrichment accelerates the consolidation of stable and task-relevant memory representations in the cortex for a novel task, and enables faster and more robust acquisition of new sequence representations.

## Introduction

Cognitive reserve may serve as a plausible explanation for why certain individuals do not experience the same level of cognitive decline otherwise manifested under neurodegenerative conditions similar in origin and extent. The complementary expression of this theory is that certain changes in brain morphology, connectivity and function, accrued from stimulating life experiences, form a cognitive reserve to help mitigate the effects of neuropathology, wherein the greater the reserve, the more serious the pathology must be in order to interfere with normal brain functioning (Stern 2002; Nunes and Silva Nunes 2021; de Rooij 2022).

While the idea of a cognitive reserve is widely acknowledged, precisely defining and measuring it has proven challenging (Cheng 2016). Environmental enrichment (EE) has been used as a rodent model of cognitive reserve and can be used to assess the neurophysiological changes resulting from complex and stimulating experiences. In fact, EE has become a standard technique for housing laboratory animals owing to the large range of benefits that it produces on health and behaviour. Engaging with complex environments, both mentally and physically, promotes the efficient use of available neural networks and the recruitment of new networks to cope with environmental demands and to adapt to brain injury (Petrosini et al. 2009). Of notable interest are the marked improvements in learning and problem-solving capacities demonstrated by enriched animals. In particular, rodents that have been exposed to complex environments with more opportunities for physical and/or social engagements have demonstrated faster learning as well as better memory retention in contextual fear conditioning (Duffy et al. 2001; Tang et al. 2001; Barbelivien et al. 2006; Gattas et al., 2022), novel object recognition (Tang et al. 2001; Bruel-Jungerman, Laroche, and Rampon 2005), the radial arm maze (Daniel, Roberts, and Dohanich 1999; Brillaud, Morillion, and de Seze 2005), and the Morris water maze task (Williams et al. 2001; Schrijver et al. 2002; Leggio et al. 2005), to name a few. EE has also been shown to mitigate the impacts of various insults to the brain in rodent models of neurodegenerative diseases and under lesions to various structures. For instance, EE has been shown to reduce spatial memory deficits in a mouse model of Alzheimer’s disease (Jankowsky et al. 2005), and to rescue a learned locomotor habit under bilateral lesions to the sensorimotor cortex (Held, Gordon, and Gentile 1985). Therefore, from a behavioural perspective, EE appears to facilitate learning mechanisms to enable the rapid acquisition of stable and resilient memories.

Consistent with these behavioural improvements in learning and memory consolidation, the neurobiological impacts of EE are manifested in plasticity mechanisms as well as anatomical and morphological changes. Enrichment has been associated with altered plasticity in the hippocampus, where larger and more persistent LTP may be induced in previously enriched animals (Duffy et al. 2001; Abraham et al. 2002). In parallel, the efficiency of synaptic transmission also saw enhancement across the major synaptic sites in the hippocampus (Green and Greenough 1986; Foster and Dumas 2001; Foster, Gagne, and Massicotte 1996). These facilitations of synaptic function are also coupled with an increase in adult neurogenesis in the dentate gyrus (Kempermann, Kuhn, and Gage 1997; Olson et al. 2006). Therefore, the behavioural improvements in learning and memory retention, especially for hippocampal-dependent tasks, are likely partly explained by the increased capacities for the hippocampus to rapidly store novel information.

In the neocortex, enrichment led to a ∼6% increase in cortical thickness (Diamond et al. 1966; Bennett, Rosenzweig, and Diamond 1969). While the overall density of neurons was reduced as a result (Turner and Greenough 1985), the volume of the cells as well as the complexity of the dendritic arborisation were expanded (Volkmar and Greenough 1972; Turner and Greenough 1985). The joint effects of EE on the hippocampus and the cortex imply that the resulting cognitive enhancements may be underpinned by more efficient communications between the two systems, hence a faster consolidation of new memories for de novo learning (cf. McClelland, McNaughton, and O’Reilly 1995). Furthermore, the higher prevalence of cortical “schemas” accumulated through enriched experiences may further facilitate the rapid assimilation of new memory traces (Tse et al. 2007; McClelland 2013). Ali et al. (2017) investigated this prospect by comparing the memory retention for a Morris water maze task between control and enriched animals that both received lesions to the reuniens and rhomboid nuclei. The latter manipulation does not impact spatial learning, but significantly shortens the retention of acquired spatial memories. Contrasted with the control group, the enriched animals exhibited better retention 25 days following acquisition, suggesting that more effective system-level consolidation had taken place.

Despite the behavioural and neurophysiological evidence implicating consolidation as a plausible mechanism underlying the benefits of EE, direct functional verification of such connections remains incomplete. In particular, whether enrichment could contribute to the rapid emergence of stable representations amongst cortical neuronal populations to support high behavioural performance is the hypothesis that the present work aims to address. In a recent study, Esteves et al. (2023) demonstrated that the formation of spatial representations in the cortex follows standard systems consolidation patterns, wherein position-correlated neuronal responses recovers following bilateral lesions to the dorsal hippocampus for a familiar environment learned prior to hippocampal damage, whereas the network’s ability to generate spatial representations for a novel environment explored following lesion was severely impaired. Thus, the hippocampus is necessary for the initial generation, but not the maintenance of cortical codes for a given environment.

Leveraging this hippocampal-dependent cortical code, this study compares the functional characteristics of cortical ensembles between enriched and exercise-matched control mice learning a virtual spatial foraging task. We also took advantage of a new EE protocol that produces substantially better improvements in cognitive function compared to standard enriched housing conditions (Gattas et al. 2022). Our results indicate that, in addition to improved behavioural performance, spatial representations became substantially more robust and more stable in enriched animals. The improved neural spatial coding in the enriched group was paralleled by an increase in the incidence of spontaneous memory retrievals (reactivation) during the early stages of learning. Taken together, these results are consistent with the notion that EE enables faster and more reliable consolidation of hippocampal-dependent memory traces into stable cortical representations.

## Results

The activity of individual neurons in superficial secondary motor cortex (M2) from enriched (N=5) and exercise-matched control (N=5) groups was recorded longitudinally using two photon calcium imaging, while animals rested and ran over a treadmill belt with 4 visuo-tactile cues across its length (Fig 1F-G). Mice were water deprived and received a 10% sucrose reward after completing a full lap. Before and after the running period (∼10 min), mice were allowed to rest for 20 min with the belt clamped (pre-Rest –R1; post-Rest – R2; Fig 1H). For data analysis purposes, only the portions of the recording during which the animals remained immobile (indicated by no input from the treadmill encoder that monitors movement) were taken into account. The same field of view (FOV) was imaged across days for one week, using cortical landmarks (blood vessels and cells) to align the FOV to that of Day 1. Consistent with our previous studies that employed calcium imaging and treadmill VR (Chang et al. 2020; Esteves et al. 2021; 2023; Chang et al. 2023), both groups had large fractions of cortical cells tuned to precise locations along the belt. Moreover, when comparing the enriched with the control group, no changes were observed in the number of cells detected across days, or the proportion of cells that passed a minimal spatial selectivity criterion (cells with place field; see Methods -Fig 2A-top). For spatially selective cells, both groups exhibited neurons with comparable spatial information (SI) and place field widths (Fig 2A-bottom).

**Figure 1:**
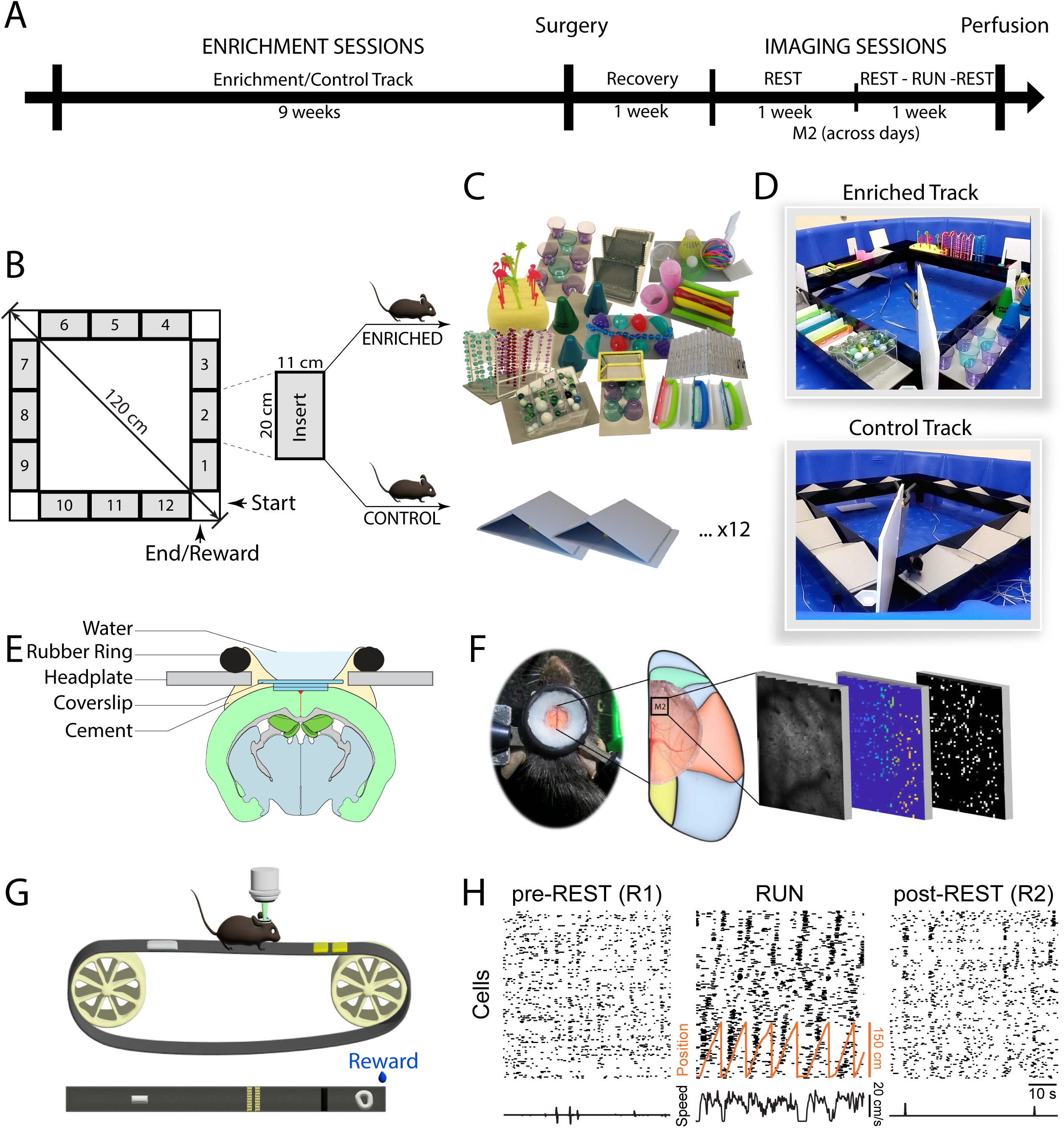
Enrichment and Imaging Paradigms. **A)** Experimental timeline. **B)** After weaning, mice started enrichment/control training. Enriched mice ran daily, 1-hour, sessions for 9 weeks on a track filled with 12 obstacles. A reward (chocolate milk) was located at the end of the track (goal). **C)** During each enrichment session, objects were added/changed/interchanged over the track to gradually increase the level of difficulty of the course (C, D-top). Control mice were exercise-matched by running a track filled with 12 ramps (C, D-bottom). **E)** Following the enrichment/control sessions, animals were implanted with a cranial window over the dorsal cortex. Animals were allowed to recover for one week and habituated in the following week to rest while head-fixed over a clamped treadmill. **F)** Two-photon Ca^2+^ imaging was used to record from the secondary motor cortex – M2 (black box). **G)** After the head-fixation/rest habituation period, animals were water restricted and imaging sessions started while the animals were allowed to rest and run/explore a 150 cm treadmill belt lined with four tactile cues. Sucrose water was delivered at the end of each lap. **H)** For a single imaging session, a 60 s segment from each imaging block is shown. Imaging sessions were divided into three blocks of 15 to 20 min each. During the first and last block (R1 and R2) the treadmill was clamped and animals were allowed to rest quietly over the belt. During the second block (run) animals explored the belt, and received reward at the end of each lap. The deconvolved ΔF/F_0_ time-course vectors of neuronal activities were sorted by their peak response positions during running. The animal’s position (red) and speed are shown below.

**Figure 2:**
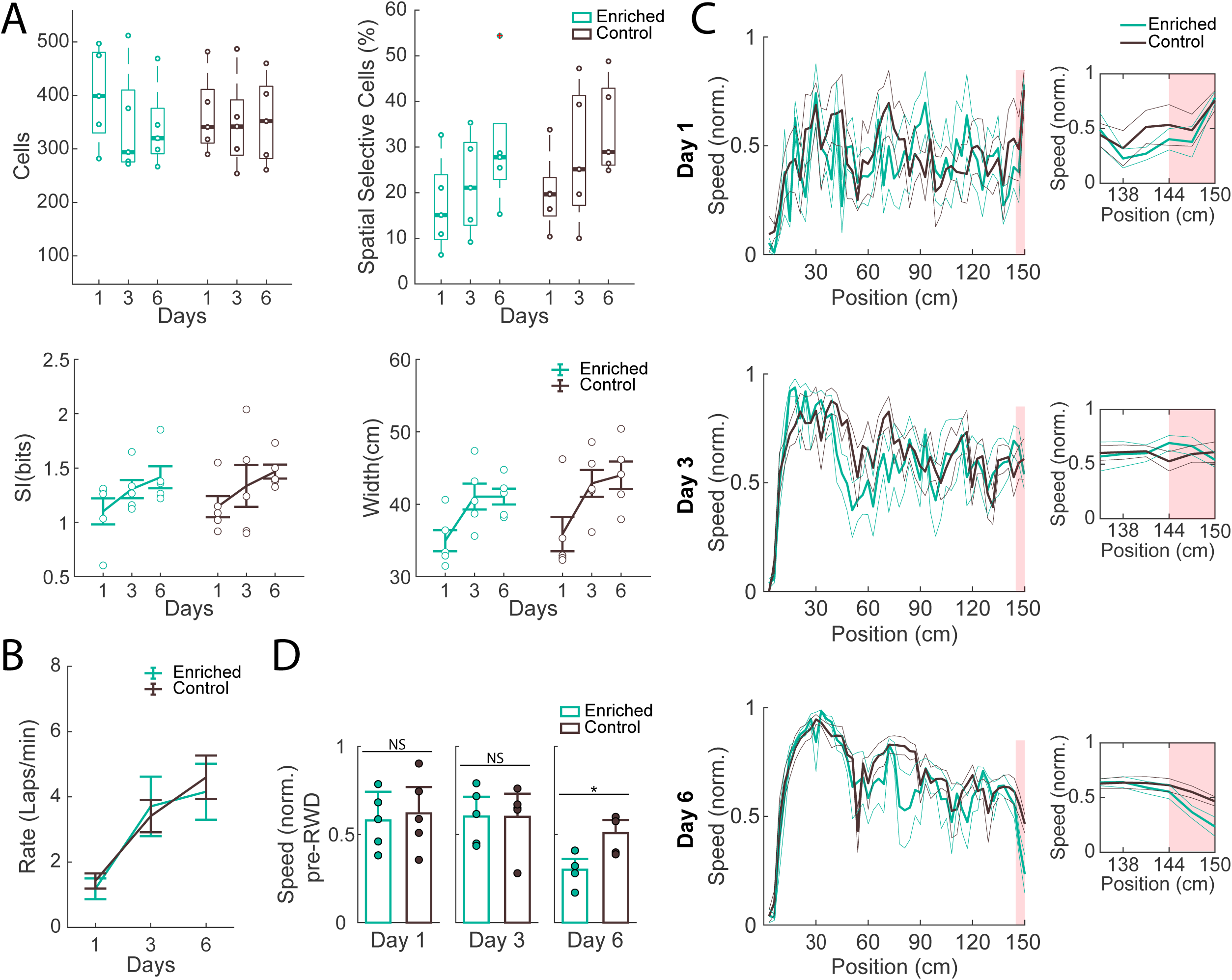
Enriched animals acquired reward anticipatory behavior earlier than controls despite co-evolving spatial tuning properties. **A-top)** Box plot of the mean number of active cells detected across mice during sessions recorded on Days 1, 3, and 6 (left); Percent of cells detected that were classified as spatially selective (using a minimal selectivity criterion) during the same days (right); Line: median; box: 25th and 75th percentiles; dots: values for individual mice; whiskers: minimum and maximum values. A-bottom) Spatial Information (left) and Place Field Width (right) of all cells classified as spatially selective on Days 1, 3, and 6. Lines: mean across all mice; dots: values for individual mice; error bars: SEM across animals No significant differences were observed between groups or group and days interaction in all measures. Significant difference between days were observed for spatially selective fraction (two-way ANOVA [F(2,24)=4.21, p=0.027)]) and Place Field Width (two-way ANOVA: [F(2,24)=7.84, p=0.02]) and marginally non-significant for Spatial Information (two-way ANOVA: [F(2,24)=3.021, p=0.068]). **B)** Running performance (laps/min) for each group during sessions 1, 3 and 6. No significant difference was found between overall running performances on the treadmill. However, statistical difference were observed across days (two-way ANOVA: [F(2,24)=10.082, p<0.01]), indicating that the animals’ performance improved with training.. **C)** Average normalized running speed as a function of position during Day 1 (top), Day 3 (middle) and Day 6 (bottom); Inset: Pre-reward speed illustrated as a function of position. Thick lines: mean across all mice; thin lines: SEM over animals **D)** Average normalized speed in the 6 cm window preceding reward onset (red shaded area on C). Error bars are SEM across mice; dots are values for individual mice; (t-test: *p=0.009). On Day 6, the enriched group exhibited a significantly greater reduction in pre-reward speed, indicating that enriched animals developed anticipatory behavior faster than the control group. For exact p values see Table 1.

**Table 1.**
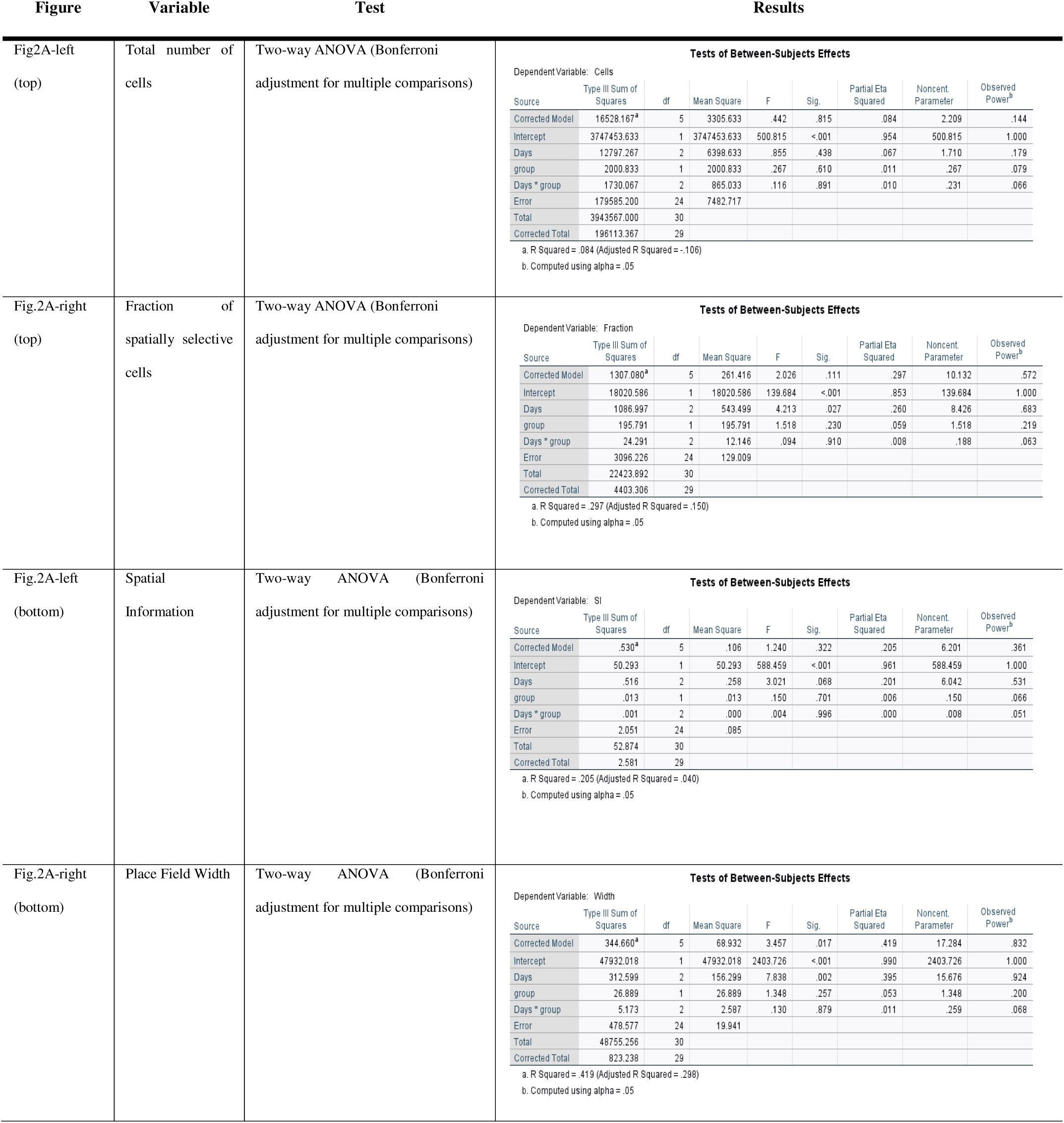
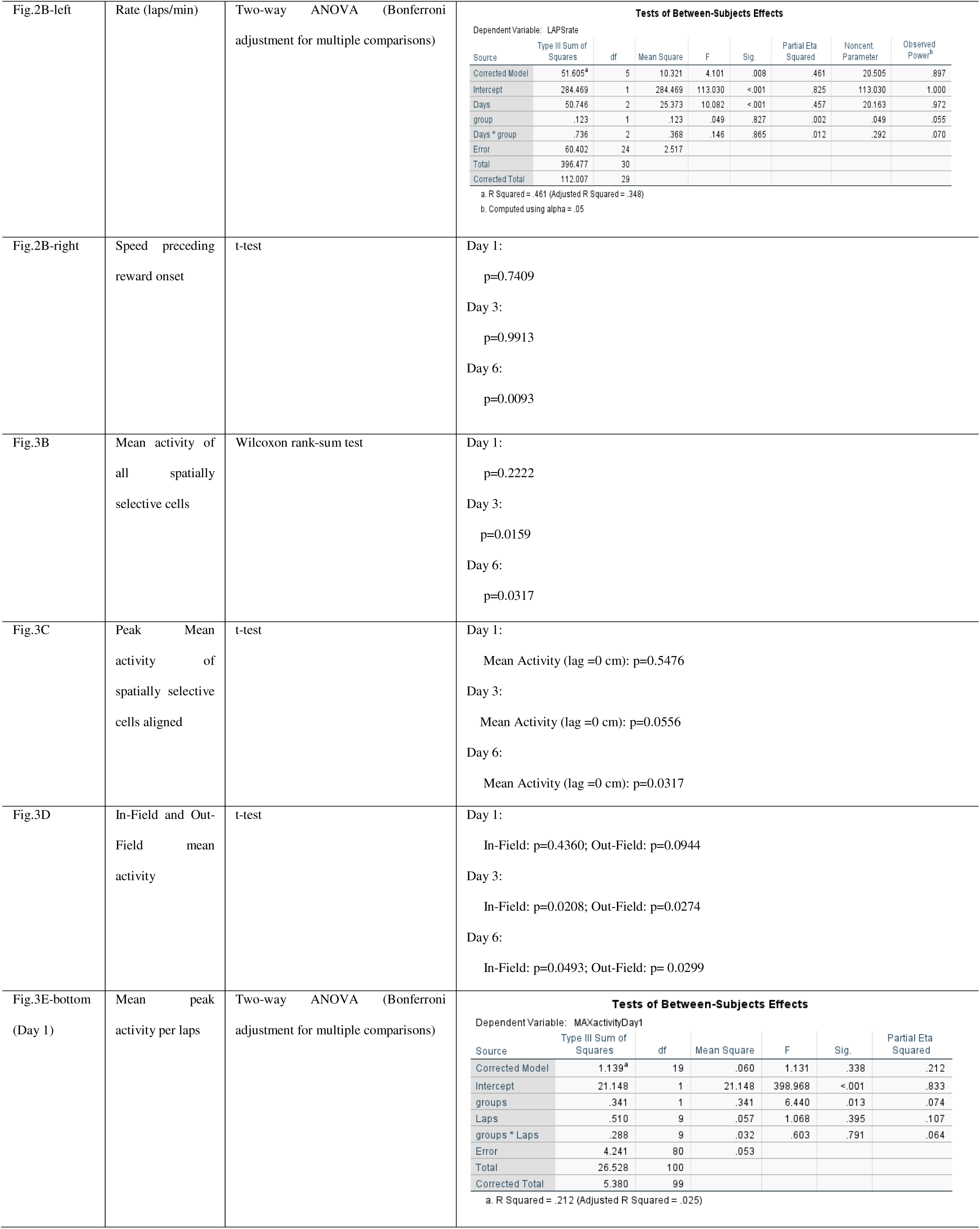
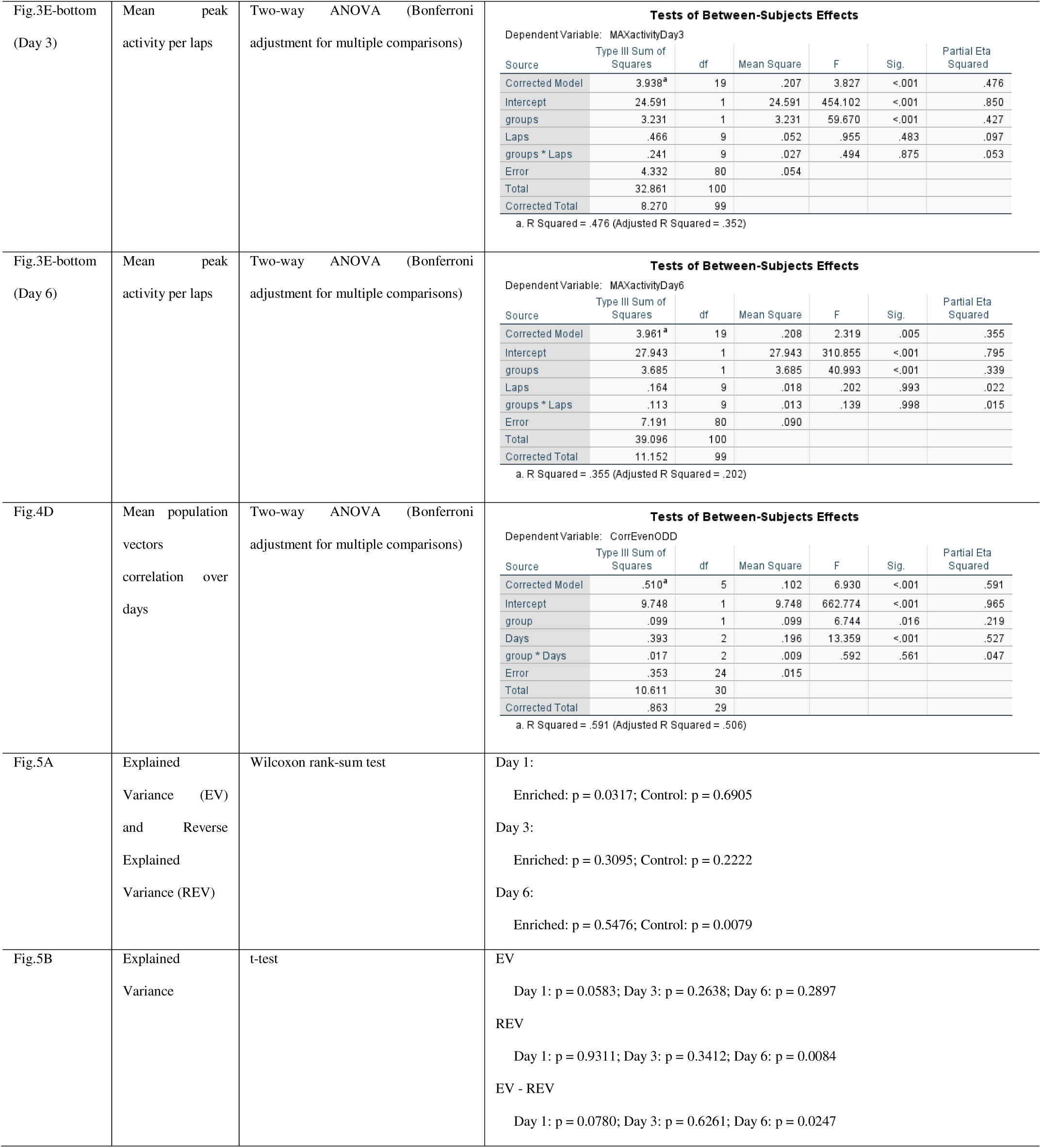
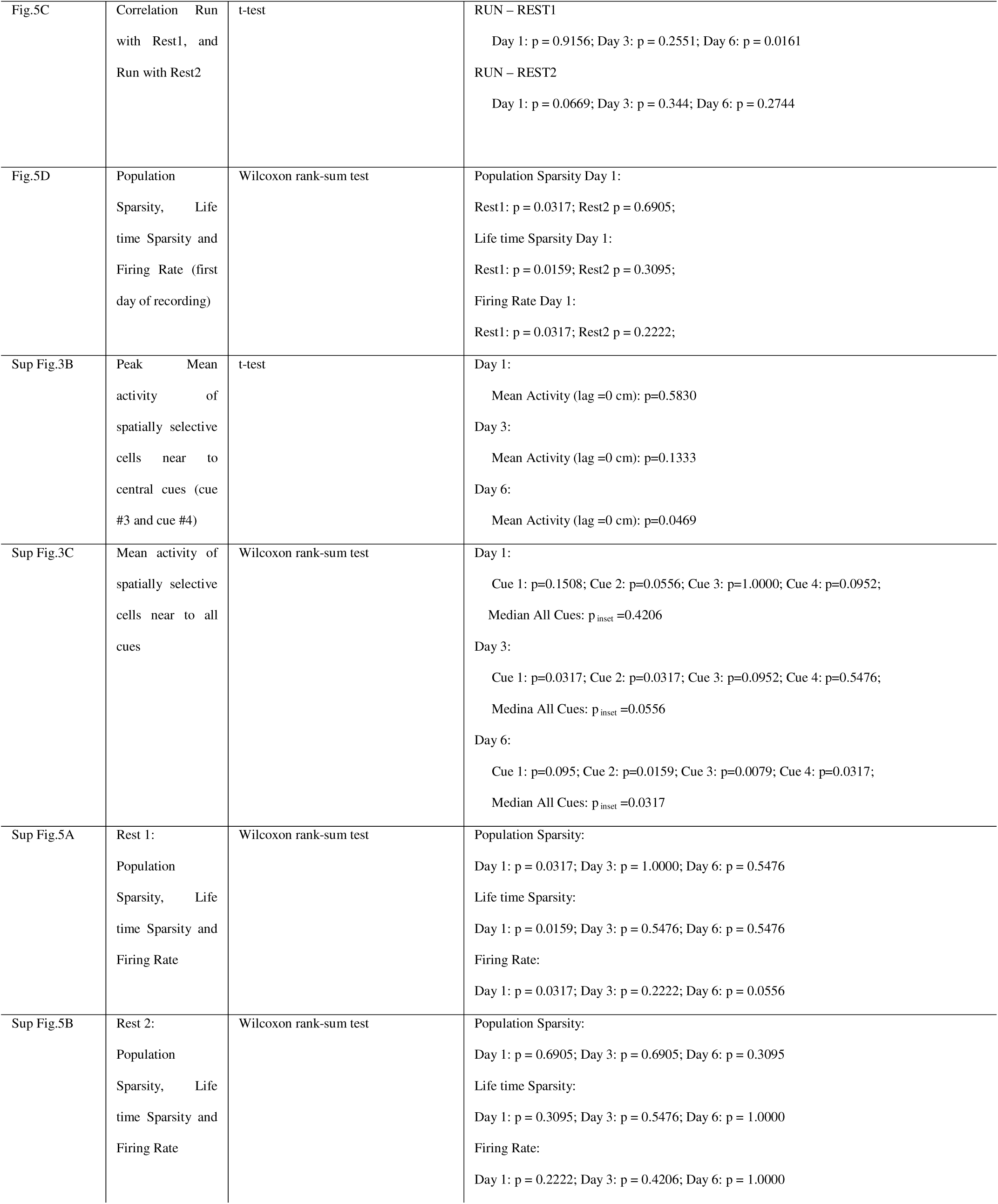
Summary of all statistics.

### Enriched animals acquired reward anticipation behaviour earlier than controls

Across sessions, animals gradually improved running performance (laps per minute), with no differences between groups (Fig 2-B). Initially, both groups exhibited similar speed profiles, which evolved comparably with learning over days (Fig 2C-top, middle). However, after only six days of performing on the treadmill, the enriched group exhibited a substantially stronger anticipatory response by reducing their speed on approach to the reward site (Fig 2C-D). The progression of the speed profile for individual animals during those days is shown in Sup. Fig 2.

### Enrichment enhanced place field activity and cue modulation throughout training

Over days, the distribution of place fields on the track (as defined by the minimal criterion) was relatively stable, with a tendency to be more concentrated near the salient cues (Fig. 3A). However, only the spatially selective neurons of the enriched animals showed an overall increase in total mean activity (ΔF/F) over days (Fig. 3B). To assess this effect more closely, we aligned all fields on their centers (Sup. Fig 3A) and averaged their mean fluorescence (Fig. 3C). In the enriched group, mean activity within the place field doubled over days, whereas no such increase was observed in the control animals (Fig. 3D-left). Meanwhile, out-of-field activity remained consistently higher in the enriched group across all days (Fig. 3D-right). A similar effect was observed when we took all neurons with place fields centered around the middle two cues in order to minimize any acceleration and deceleration effects (Sup. Fig 3B). The overall mean activity near environmental cues also increased over days but only for the enriched group (Sup. Fig 3C). We also examined how place fields evolved over the first ten laps of each session (Fig. 3E-top). Spatially selective cells in both groups remains relatively stable across laps during the three days (Fig. 3E-bottom). These findings suggest that the enriched group exhibits neurons whose activity increases with training. In contrast, the control group shows lower out -field activity compared to the enriched group, and place field activity does not increase over time.

**Figure 3:**
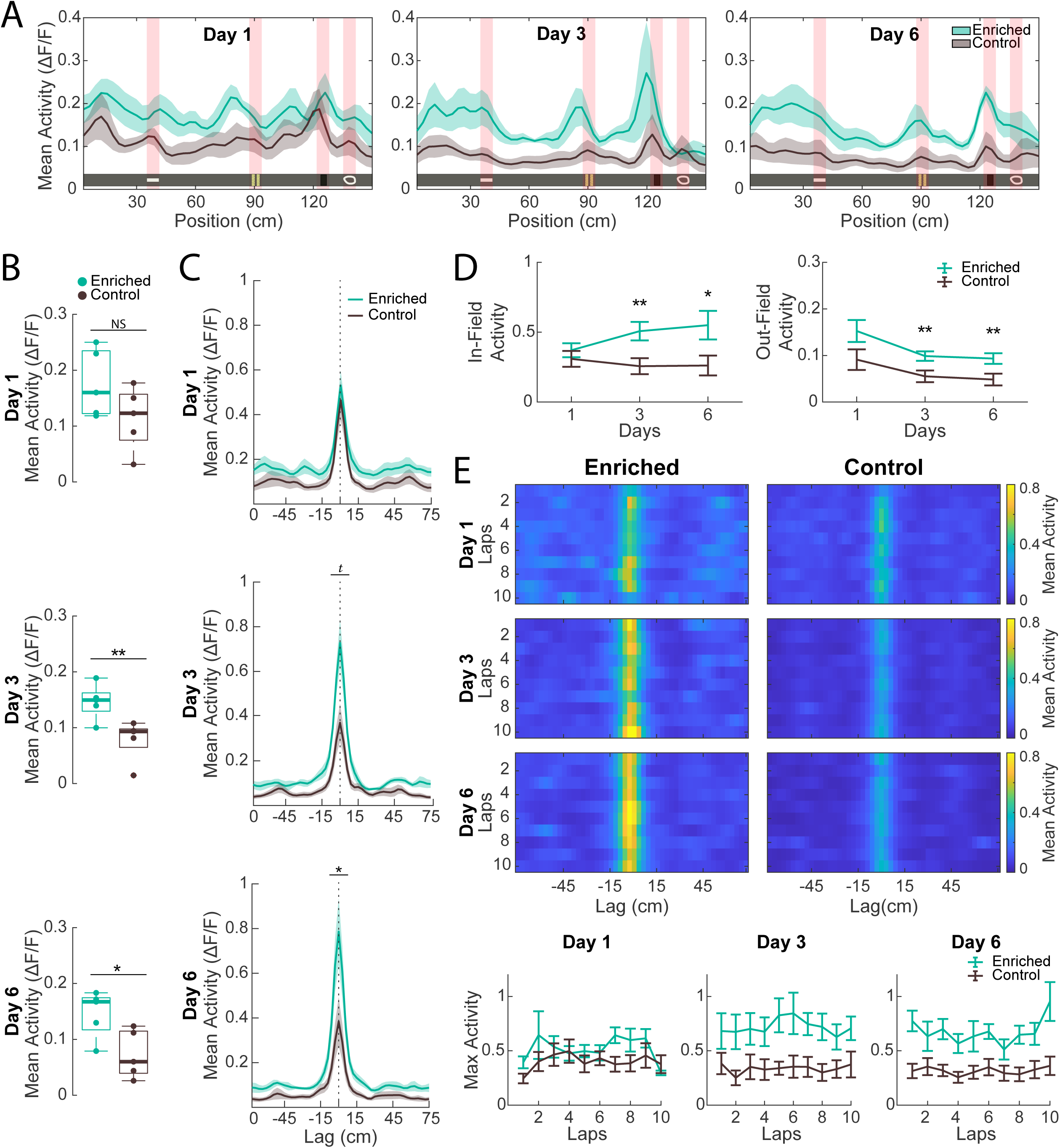
Place fields in enriched mice showed progressively increased activity over recording sessions. **A)** Trial-averaged activity as a function of location (non-normalized) of all cells that presented a place field on Days 1, 3, and 6. Green and gray shaded area are SEM over animals. **B)** Box plot of the mean activity of same cells shown in A. Following training, enriched group exhibited significant higher ΔF/F_0_ (Wilcoxon rank sum test: NS=not significant; **p=0.0159; *p=0.0317). Line: median; box: 25th and 75th percentiles; dots: values for individual mice; whiskers: minimum and maximum values. **C)** Cells with place fields identified on Days 1, 3, and 6. Place fields were aligned by their centre and averaged across groups. Green and gray shaded area are SEM over animals (same cells showed in Sup Fig 3A). In the enriched group, place field intensity doubled over time, whereas no such increase was observed in the control group (t-test: ^t^p=0.0556; *p=0.0317). **D)** Average in-field activity (lag: -6cm to +6cm) and out-of-field activity (lag: -75 to -15 and +15 to +75) for all cells with place fields on Days 1, 3, and 6, showed in B. Error bars are SEM across mice. Significant group differences were observed exclusively on Days 3 and 6 for both in-field and out-of-field activities. (t-test: *p<0.05 **p<0.03). E-top) Trial-averaged activity maps of place cells across the first 10 laps on Days 1, 3, and 6. E–bottom) Mean peak activity (lag = 0) as a function of the first 10 laps on Days 1, 3 and 6. Error bars represent SEM across animals. We observed a main group effect for all days (two-way ANOVA Day1: [F(1,80)=6.440, p=0.013]; Day3: [F(1,80)=59.670, p<0.01]; Day6: [F(1,80)=40.993, p<0.01]). Overall, with training, the enriched group exhibited a general increase in in-field neural activity. For exact p values see Table 1.

### Place fields of enriched mice became more pronounced and stable across sessions

Next, we evaluated the stability of spatial representations over days. We began by identifying spatially selective cells on Day 4 that were active on at least one of the subsequent days (Sup. Fig 4), and we noticed that the representation of environmental cues appeared more stable across days for the enriched group (Fig 4A). When observing the average mean activity over locations (Fig 4B), we noticed that cells with place fields near cue locations presented an overall higher activity. We then computed the Pearson correlation matrix of the neural population codes for location detected on day 4 with the subsequent two days (Fig. 4C). The diagonal of these maps for Day 5 and 6 represents the similarity in population vectors at the same locations across days. For Day 4, we computed the correlation between even and odd trials in order to indicate the effects of trial by trial variability. The average correlation coefficient of the diagonal shows that, while there was representational drift in both groups, the effect was significantly more pronounced in the control group (Fig. 4D). Overall, across all days, the mean correlation between population vectors for the control group shows lower similarity (i.e., lower stability and more drift) than for the enriched group (Fig. 4D-E). Altogether, these findings indicate that the enriched group developed a neural representation of the virtual environment that was more robust and more stable over time.

**Figure 4:**
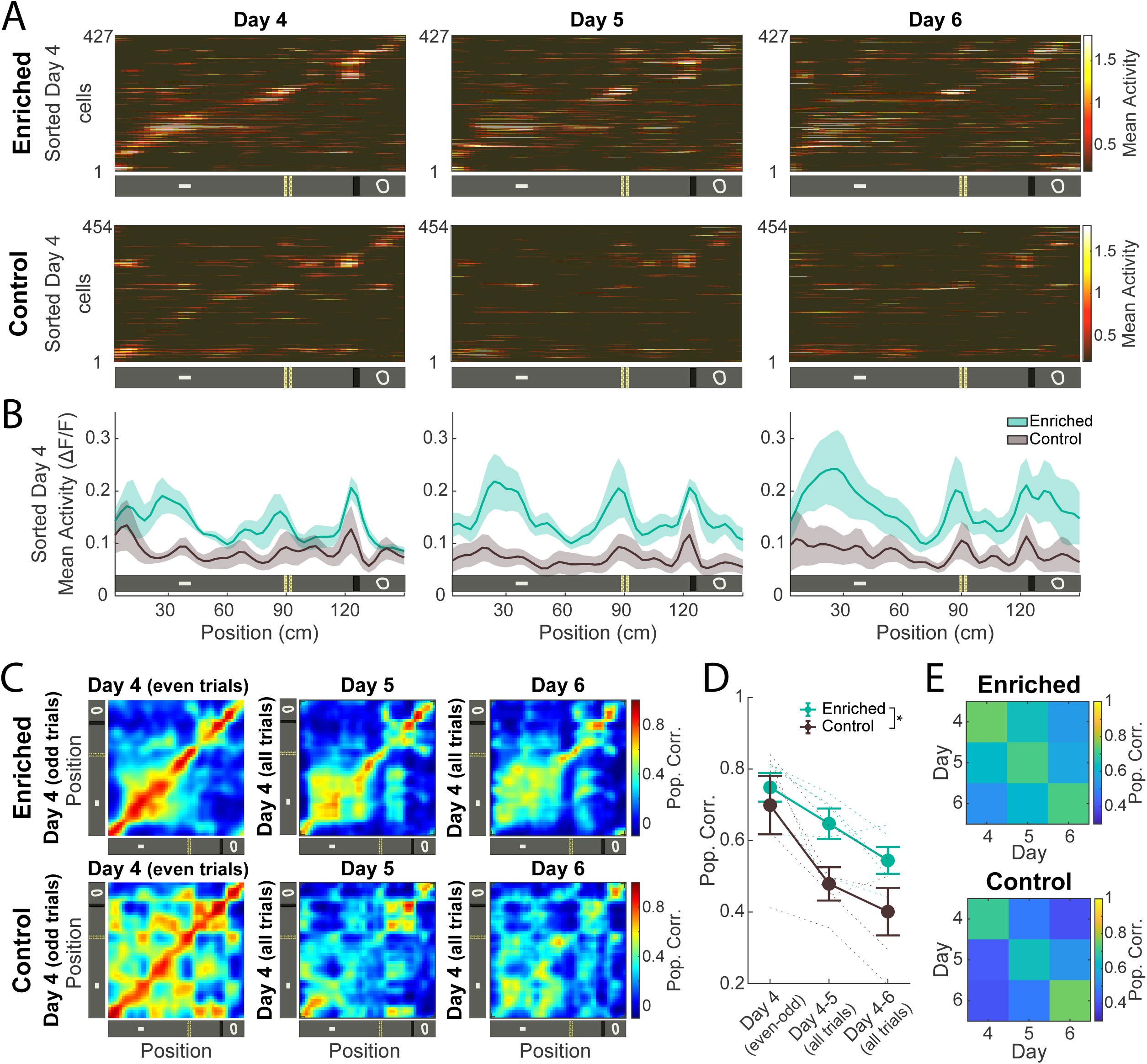
Enrichment improved the stability of cortical spatial representations across days. **A)** Trial-averaged activity map as a function of location (non-normalized). Cells are sorted by Day 4 place field locations. Only spatially selective cells detected on Day 4 are shown on Day 5 or Day 6. Data pooled across all animals of each group. **B)** Mean activity as a function of position for neurons shown in A. Shaded area are SEM over animals. **C)** Pearson correlation matrices of population vectors for three consecutive days, using even and odd trials for Day 4 (same cells showed in A). **D)** Mean population vectors correlation over days (diagonal of matrices shown in E). Error bars are SEM over animals. There was a significant effect over groups (two-way ANOVA: [F(1,24)=6.744, p=0.016]) and days (two-way ANOVA: [F(2,24)=13.359, p<0.001]) without significant group and days interaction. **E)** Mean population vectors correlation between spatially-selective cells on a reference day (y-axis) that remained active on the comparison day (x-axis). The enriched group expressed neurons with higher stability, indicated by cells with higher correlation during the subsequent day. For exact p values see Table 1.

### Enriched animals exhibited early-stage cortical memory trace reactivation

We used explained variance (EV; Kudrimoti, Barnes, and McNaughton 1999) to quantify the fraction of the variance in neural population activity during RUN that could be explained by the post-REST activity, while accounting for any intrinsic/confounding correlations between pre-REST and RUN (Tatsuno, Lipa, and McNaughton 2006). As a control, we compared the EV against the reverse-EV (REV; Pennartz et al. 2004). The latter is used to estimate similarity between Run and pre-Rest periods (R1). We found that, during the first day, EV was significantly higher than the REV only for the enriched group (Fig 5A-left), indicating that the new experience of learning how to navigate the virtual environment was being reactivated in the secondary motor cortex (M2) during the post-task rest. For the control animals, the magnitude of reactivation increased with experience and was only evident on Day 6 (Fig 5A-right). In contrast, across the learning days, EV and REV indicate that neural activity in enriched animals during the Run period became less explained by the post-Rest session and better explained by the pre-Rest session (Fig 5B-left, middle). We also found that, while the reactivation strength (EV-REV) of the control group increased with experience, there was a decay of the reactivation value across sessions for the enriched group (Fig 5B-right). To evaluate the degree of similarity between Run and Rest sessions, we computed the Pearson correlation between cell-pairs across Run-R1 and Run-R2. These results show that while exposure to the task increased the similarity between pre-Rest and Run for the enriched groups, the opposite effect happened with the control group (Fig 5C-top). Moreover, the Run session of the enriched animals on Day 1 had a tendency to show a greater degree of resemblance to the post-Rest session, and this similarity decreased with learning and time (Fig 5C-bottom). As a result, following the enriched animals’ the initial exposure to the virtual environment, there was a greater degree of similarity between Run and R1 in terms of correlation structure, and R1 accounts for a larger portion of the variance in the Run session. These findings suggest that, after a given amount of exposure to the task, cortical neurons in the enriched groups can predict the neural activity of the Run session during the pre-Rest period.

**Figure 5:**
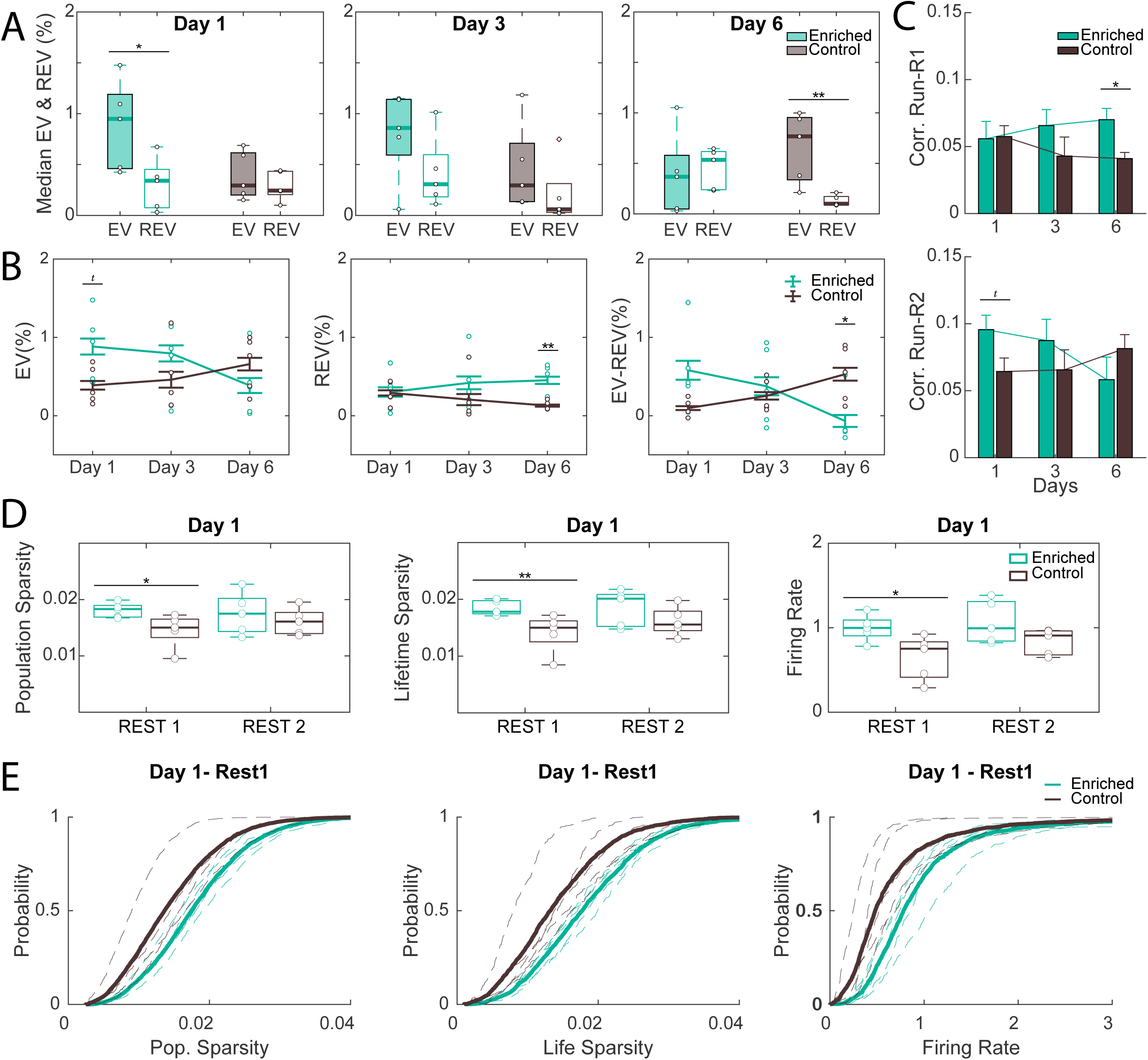
Enriched animals exhibited early-stage cortical memory trace reactivation. **A)** Median explained variance (EV) and reverse explained (REV) encoded by all cells during sessions recorded on Days 1, 3, and 6. Significant reactivation was observed during Day 1 for the enriched group (EV > REV), and this difference reduced over time. For the control group reactivation was significant only on Day 6 (Wilcoxon rank sum test: *p=0.0317; **p=0.0079). Line: median; box: 25th and 75th percentiles; dots: values for individual mice; whiskers: minimum and maximum values. **B)** Percentage of explained variance, reverse explained variance, and EV-REV (measure of post-Rest reactivation) are shown for sessions on Day 1, 3, and 6 (t-test: ^t^p=0.0583; *p=0.0247; **p=0.0084). Error bars are SEM across mice. The enriched group showed a higher EV on Day 1 and a significantly higher REV on Day 6. **C)** Average Pearson correlation coefficient between cell-pairs for Rest1 and Run sessions (upper), and Rest2 and Run sessions (bottom) during Days 1, 3, and 6 (t-test: ^t^p=0.0669; *p=0.0161). Error bars are SEM across mice. Run sessions of the enriched group tended to present a higher degree of similarity with Rest 2 only during the first day of training. On Day 6, after learning the virtual environment, both groups showed similar Rest2-Run correlation values. Moreover, the enriched group exhibited population activity of Run with a greater degree of resemblance with Rest1 (pre-Rest). **D)** Boxplots of population sparsity, lifetime sparsity, and firing rate encoded by all cells detected during the first imaging session (Day 1). Enriched animals present a higher Firing Rate, a lower lifetime sparsity (higher index) only before the first running session (Wilcoxon rank sum test: **p=0.0159; *p=0.0317). **E)** Cumulative Distribution of same data showed in D. -dashed lines: values for individual mice; thick lines: mean across all mice. Box Plot - Line: median; box: 25th and 75th percentiles; dots: values for individual mice; whiskers: minimum and maximum values. For exact p values see Table 1.

The population sparsity, lifetime sparsity, and firing rates levels were not statistically different between the two groups across training days, with the notable exception of the initial resting state (R1-Day1) where the enriched group displayed both statistically higher firing rates and lower sparsity (more distributed) compared to the control group (Fig 5D-E; Sup. Fig 5).

In summary, enriched group showed stronger memory trace reactivation when first exposed to a virtual environment. Subsequently, the neural activity of enriched animals during the task became more similar with pre-task activity, suggesting anticipation of the experience, while the control group showed the opposite trend. Despite these differences, the firing rates and sparsity of neuronal populations were comparable between the groups throughout training, with the exception of the first pre-exposure rest period, during which enriched animals had higher firing rates and more distributed activity.

## Discussion

Prior exposure to an enriched environment accelerates the rate at which mice acquire stable and accurate cortical representations of locations in a virtual environment. Over days of exposure to the foraging task, neurons in the secondary motor cortex of the enriched group exhibited an increase in place field strength and a smaller degree of representational drift/higher stability compared to the control group, supporting stable correlates of firing and position. This fast stabilization of the neuronal sequences was paralleled by an increase in the rates of reactivation in the enriched group on the first day of exposure, along with a potential prediction of the upcoming experience. In contrast, the opposite trend was true of the control animals, where reactivation gradually increased with repeated exposures. These functional observations are commensurate with the behavioural anticipation of upcoming rewards, which enriched animals began to exhibit within a week of training. Taken together, these results suggest that environmental enrichment accelerates the rate of consolidation and stabilization of a hippocampal-dependent trace into the cortical network.

Behaviourally, it has been shown that environmental enrichment improves spatial learning and the ability to predict reward locations (Dhanushkodi et al. 2007; De Bartolo et al. 2008; Templer, Wise, and Heimer-McGinn 2019). Under head-fixed preparations, such learning performance can be effectively quantified through anticipatory behaviors, such as licking or decelerating before reward delivery (Burke et al. 2005; Jurjut et al. 2017; Garrett et al. 2020; Garner and Keller 2022). In particular, it has been shown that licking and slowing down before the reward delivery point develop after 5-10 training sessions (Sato et al. 2020; Esteves et al. 2023; Issa et al. 2024). Here, animals that underwent enrichment began exhibiting deceleration on approach to reward at earlier stages of training than controls, suggesting that prior enrichment increases the rate of novel spatial learning.

Crucially, the faster behavioural expression of learning was associated with the gradual stabilization of neuronal representations relevant to the task. In general, cortical representations that are correlated with behavioural parameters tend to emerge and fluctuate with experience until a level of stability is reached with expertise (see for review: Rule, O’Leary, and Harvey 2019; Driscoll, Duncker, and Harvey 2022). For spatial representations, in the hippocampus, stability over time varies widely by region (Lu et al. 2015; Hainmueller and Bartos 2018), although a baseline level of representational drift is always present despite strong familiarity (Ziv et al. 2013; Rubin et al. 2015; Driscoll, Duncker, and Harvey 2022). In contrast, neural activity in the cortex that correlates with sensation, movement, and cognition appears to remain more stable across time (Driscoll et al. 2017; Demchuk 2023). Specifically, previous reports suggest that this stability may be reached following ∼10 days of exposures to a given environment (Esteves et al. 2023). Of notable interest is a recent study in which it was reported that spatial representations in the medial prefrontal cortex became markedly more stable following learning of an odour-guided choice task, suggesting that the stabilization of cortical spatial representations is strongly correlated with learning (Muysers et al. 2024). Therefore, the greater stability of cortical representations exhibited by environmentally-enriched animals over days, compared to their exercise-matched counterparts, is indicative of a faster reshaping of neuronal representations to support stable learned behaviours.

How enrichment improves learning and the stabilization of cortical representations may have to do with the rate and readiness at which novel experiences are consolidated. It has been shown previously that, in a flavor-place paired association task, rats were able to quickly acquire a new set of associations when the new associates were sufficiently similar to the previously learned ones (Tse et al. 2007). In contrast to schema-inconsistent associations, these new, schema-consistent associations were retained following hippocampal lesions performed 48 h after learning, but not after 3 h. Therefore, it would appear that the availability of a prior schema accelerates de novo learning by accelerating the rate of consolidation. The task employed in the present study is known to be hippocampal-dependent, wherein the formation of the position-correlated sequences in cortical areas is severely impaired in mice that received bilateral lesions to the dorsal hippocampus (Mao et al. 2018; Esteves et al. 2021; 2023). It is therefore possible that the heightened stability of cortical representations in enriched animals reflects an increased capacity for hippocampal information to be quickly integrated into cortical stores owing to the existence of more diverse priors accrued from enriched experiences.

Further corroborating the idea that faster rates of consolidation may underlie the learning benefits associated with environmental enrichment are the different patterns of reactivation observed in enriched and control groups. Reactivation, the spontaneous reinstatement of activity patterns associated with a previous behavioural experience, is a strong correlate of learning and consolidation of novel memory traces. The prevalence of reactivation during sleep has been shown to predict subsequent performance on a spatial memory task (Dupret et al. 2010; Singer et al. 2013), whereas artificial disruption of reactivation impaired retention of spatial memories (Ego-Stengel and Wilson 2010). In the present study, reactivation in the motor cortex was evident in enriched animals on the very first day of exposure to a novel virtual environment, while it was not until the sixth day of exposure that reactivation was detected in control animals. These results, therefore, suggest that cortical consolidation of neural representations occurs early-on with enrichment, while this process is more gradual in naive/impoverished animals.

One study has shown that, in the hippocampus, reactivation persisted for up to 10 h following exposure to a novel environment, but the incidence of reactivation gradually decreased down to a few minutes with repeated exposures and familiarity (Giri et al. 2019). From these observations, one could infer that the rate of learning and consolidation should be tied to the rate of decay in reactivation rather than the onset time of reactivation as presently reported. This discrepancy may be explained by the differential involvements of the cortex and the hippocampus over the time course of consolidation. In fact, whereas the hippocampus is theorized to act as a fast-learning memory system, consolidation is believed to occur gradually in the cortex (Marr 1971; McClelland, McNaughton, and O’Reilly 1995; Saxena and McNaughton 2024). Similarly, the type of information and representations reactivated by the cortex and the hippocampus appear to be different, with the hippocampus concerned with global contextual information and the cortex retrieving contents pertaining to sensory and cognitive features (Wilson and McNaughton 1994; Peyrache et al. 2009; Chang et al. 2020; 2023). Therefore, the prevalence of reactivation in the cortex could possibly reflect the gradual consolidation and formation of stable representations, where task-relevant feature sets initially ‘indexed’ by the hippocampus over time become associated and independently retrievable within cortical sites (Teyler and DiScenna 1986; Teyler and Rudy 2007; McNaughton 2010).

Taken together, the findings indicate specific functional changes in the cortex arising from prior exposures to an enriched environment. Alongside the faster emergence of anticipatory behaviors in animals from enriched group, these results suggest that novel experiences can be more readily and rapidly consolidated into existing cortical representations to promote learning. Using a hippocampus-dependent task, we demonstrated that novel representations of the belt position reach stability earlier in the cortex following enrichment, and reactivation occurs more strongly upon the first exposure to the novel task. A better understanding of the brain alterations that occur during enrichment could lead to the development of novel approaches to preserve or recover cognitive function in people living with neurodegenerative diseases.

## Author contributions

Conceptualization: BLM, IE; Experiments: ARN, IE; Data analysis and interpretation: BLM, HC, IE; Writing: BLM, HC, IE. All authors revised the manuscript.

## Funding

This work was supported by the National Institutes of Health, USA (NIH) [grant #NS121764 (B.L.M); #RF1NS132041 (BLM); #MH125557 (BLM)], and by the Natural Sciences and Engineering Research Council of Canada (NSERC) [grant #1631465 (B.L.M)], and by the Canada Graduate Scholarships for Doctoral Program (NSERC CGS-D; HC).

## Acknowledgement

This research was enabled in part by support provided by WestGrid (www.westgrid.ca) and Compute Canada (www.computecanada.ca). We thank Isabelle Gauthier, Di Shao, animal care staff and all other members of the Animal Welfare Committee at the University of Lethbridge for enabling experiments on animals. We thank Bailey Porter and Nicolas Villamil for training the animals. We thank Yagika Kaushik, Rui Pais, Ritwik Das, Rajat Saxena, Rob Bain, Michael Eckert, and Janina Ferbinteanu for their insightful and valuable comments. Further, we thank Amanda Mauthe-Kaddoura, Jennifer Tarnowsky, Maurice Needham, Andrew McNaughton, Valerie Lapointe and Karim Ali for providing technical and/or logistics support, and Dr. M. Mohajerani for supervision/administration of microscope and animal breeding facilities.

## Financial Disclosures

None of the authors have any potential or actual financial interests or conflict of interest.

## Figures Captions

**Sup. Figure 1:**
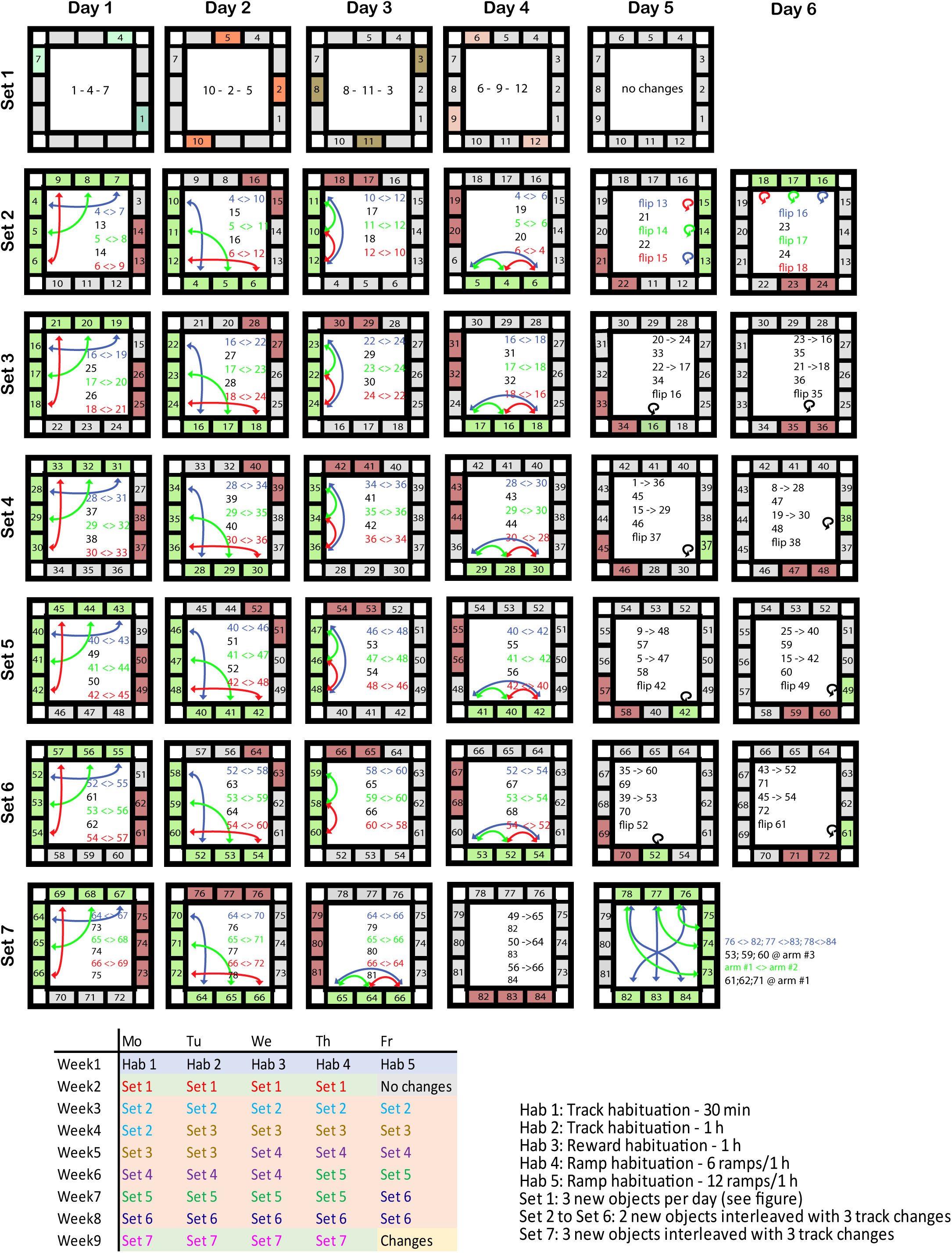
Overview of the Enrichment Training Protocol. The enrichment track was filled with a total of 12 obstacles. During each hour-long session, five to six course manipulations were made at 10-12 min intervals. Depending on the session, the manipulation included two to three new insert replacements (shown in red) interleaved with three insert rearrangements (rotating or swapping, marked by arrows). After nine weeks of training, the enriched animals had gone through 84 different obstacles, and the track was renewed every five to six sessions with a new set of 12 objects (Set1 to Set7).

**Sup. Figure 2:**
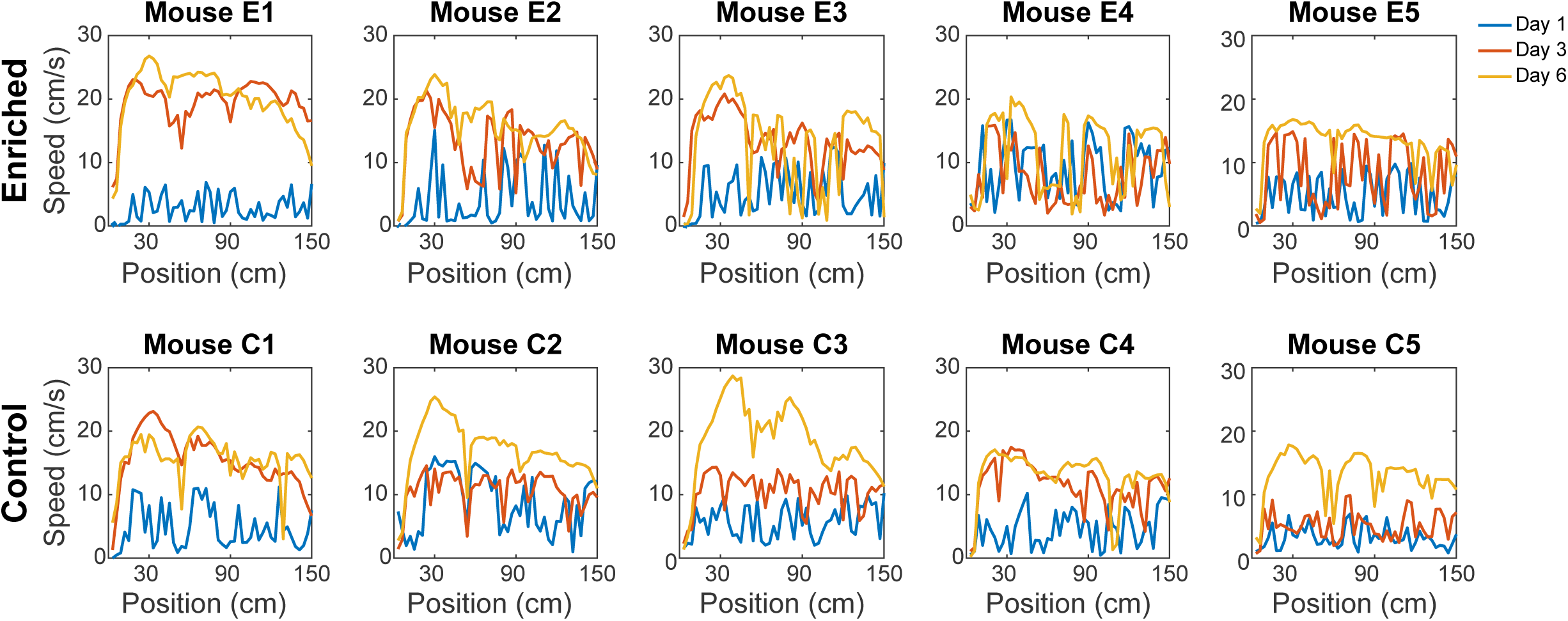
Behavioural changes across sessions for each individual animal, as they learned to run on the treadmill. Average running speed as a function of position for all animals during running sessions performed on: Day 1 (blue), Day 3 (red) and Day 6 (yellow).

**Sup. Figure 3:**
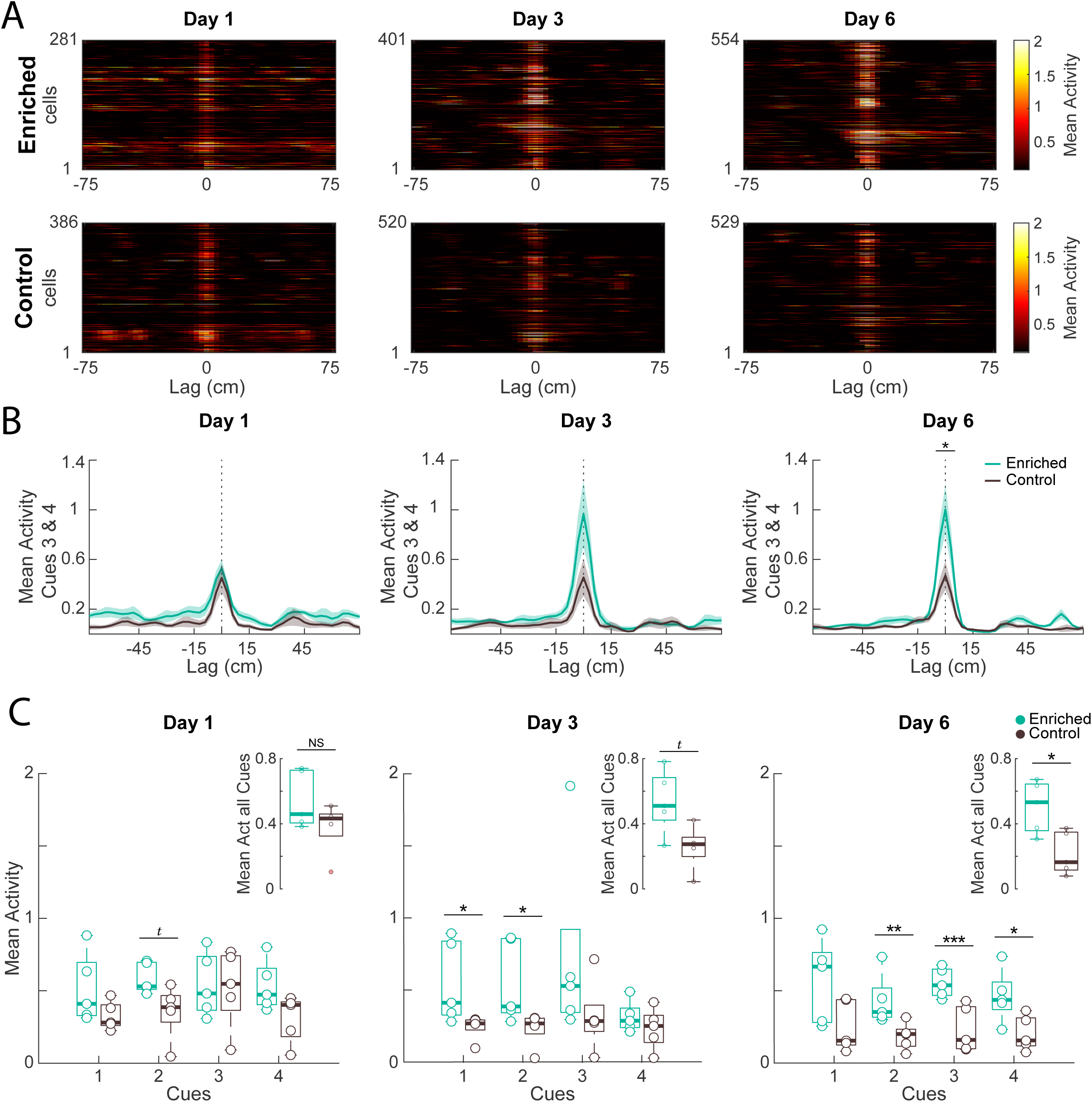
Enriched group presented an increased mean activity preceding each cue as they learned to run on a treadmill. **A)** Trial-averaged activity map as a function of location (non-normalized) for all the cells that presented a place field on Day 1, Day 3, and Day6, with peak aligned in the center (Lag = 0cm). Data pooled across all animals of each group. **B)** Cells with place fields located near the two central cues on Day 1, 3, and 6, fields were center-aligned and averaged across groups. Green and gray shaded area are SEM over animals (t-test: *p=0.0469). **C)** Boxplot of the mean activity close to each cue. Inset: Boxplot of the mean activity over all cues. Following training, enriched group exhibit an increased mean ΔF/F_0_ activity preceding the great majority of the cues location (Wilcoxon rank sum test: NS=not significant; ^t^p=0.055; *p=0.031; **p=0.015; ***p=0.007). Line: median; box: 25th and 75th percentiles; dots: values for individual mice during one session; whiskers: minimum and maximum values; + signs: outliers. These data indicate that, over days, the enriched animals developed significantly greater ΔF/F₀ near the environmental landmarks, whereas the control mice did not. For exact p values see Table 1.

**Sup. Figure 4:**
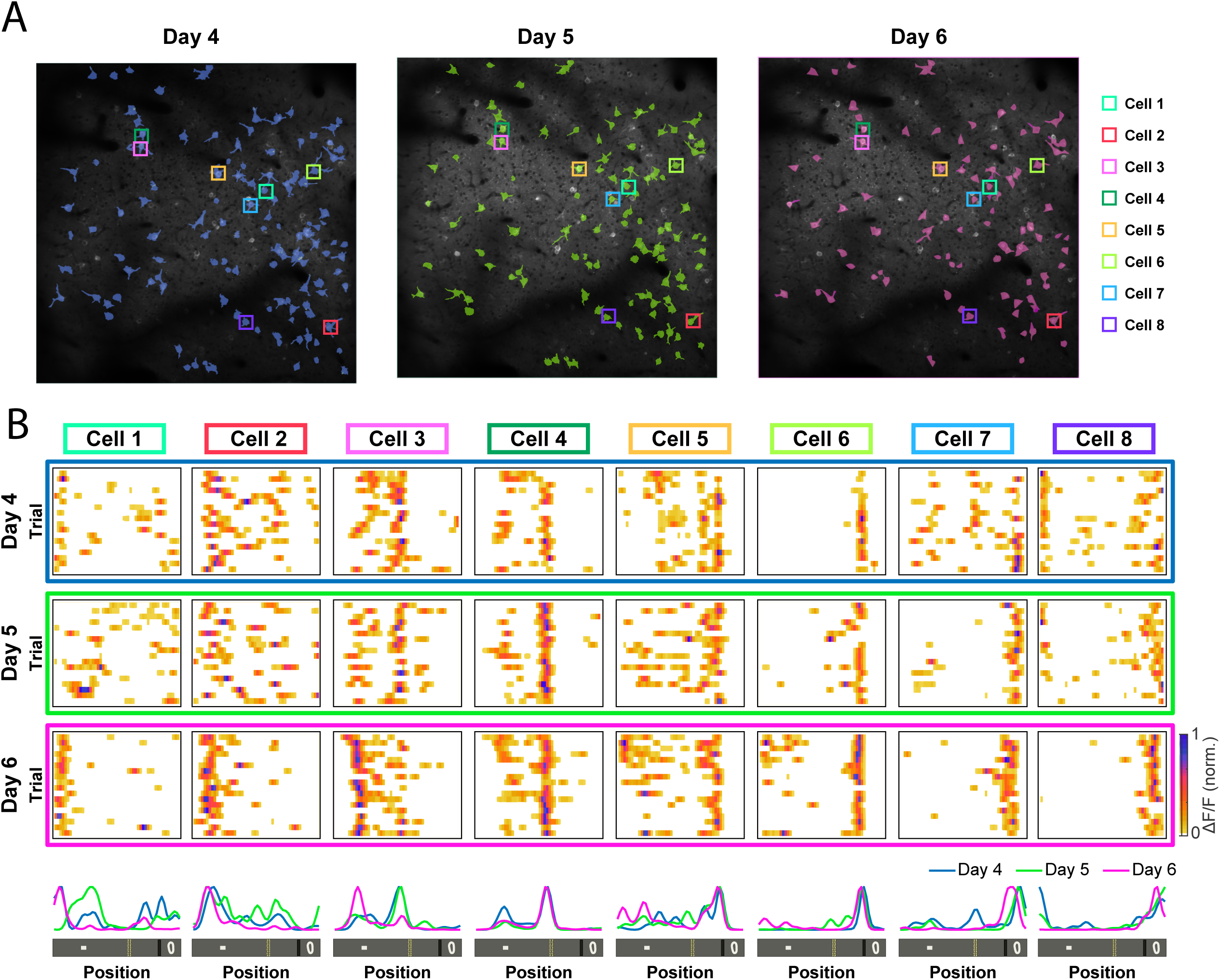
Illustrative example of M2 cells from an enriched animal imaged across three consecutives sessions. **A)** Mean imaging planes on Day 4, 5, and 6. Overlapped ROIs indicate spatially selective cells detected on Day 4 (blue) that were tracked on Day 5 (green) and on Day 6 (magenta). **B-top)** Trial by trial calcium activity (ΔF/F normalized) over position for the eight cells outlined in A by a box on corresponding days. **B-bottom)** Average tuning curve (mean calcium activity normalized) as a function of position for all eight neurons during the three consecutive days.

**Sup. Figure 5:**
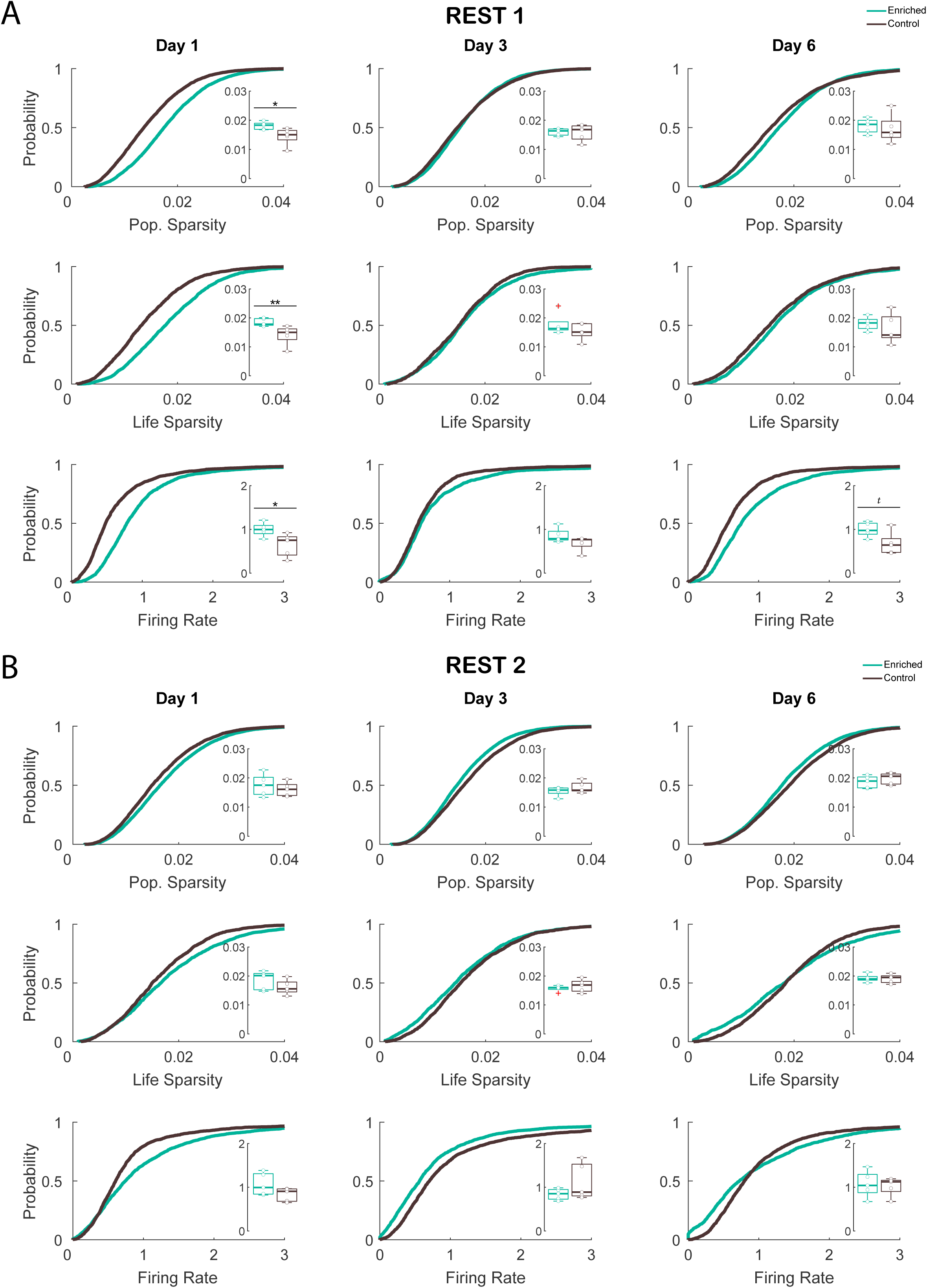
Enriched animals showed neurons with reduced lifetime sparseness and a higher firing rate before they learned to run on a treadmill. **A)** Cumulative distribution of population sparsity (top), lifetime sparsity (middle), and firing rate (bottom) encoded by all cells found on Day 1, Day 3, and Day 6 during Rest 1. **B)** Same as **A** for Rest 2. Inset: Boxplots of the population sparsity, lifetime sparsity, and firing rate for the same sessions (Wilcoxon rank sum test: **p=0.0159; *p=0.0317; ^t^p=0.0556). Line: median; box: 25th and 75th percentiles; dots: values for individual mice during one session; whiskers: minimum and maximum values; + signs: outliers. For exact p values see Table 1.

## Methods

### Experimental Design

Animals were randomly split into enriched and control groups and were trained for 9 weeks to run on an enriched or control track, respectively. Both groups underwent cranial window implantation above the secondary motor cortex (M2) and, after one week of recovery, animals were head-restrained and habituated to rest over a clamped treadmill (one hour/day during one week). Following this habituation period, mice were water restricted, the treadmill was unclamped, and they began running for reward over a 150 cm treadmill belt that had four tactile cues spaced at irregular intervals. The animals had not previously experienced treadmill running and so the treadmill was entirely novel to them. Two-photon calcium imaging was carried out while animals learned to run over the treadmill (Fig 1A). We monitored the same neurons across days over the course of a week to assess the impact of enrichment on the consolidation of spatial memory traces.

### Animals

Ten 28-day old male transgenic Thy1-GCaMP6s mice were employed in this study. The animals were randomly divided between groups and housed in standard rodent cages in pairs (one enriched and one control per cage) with food and water available *ad libitum* until the beginning of treadmill running sessions. The housing room was kept at 2411 under a 12h light/dark cycle, with lights on at 7:30 AM. All experiments were carried out during the light cycle using protocols approved by the Animal Welfare Committee of the University of Lethbridge. All procedures were conducted in compliance with the guidelines established by the Canadian Council on Animal Care.

### Enrichment Track

The enrichment protocol was designed essentially according to Gattas, et al. (2022). Mice started the enrichment training at the age of 28 days, for five days a week over nine weeks. The training session of the enriched group was one hour long, during which the mice ran a custom-made square-shaped track gradually filled with 12 obstacles that became more complex over the course of the training sessions. The control group ran for one hour on an identical track filled with 12 ramps (Fig 1B-C). For both groups, the tracks, which were made of black acrylic, had 7 cm surrounding walls to keep the animals within the course. A wall divider was placed between the start and end points to ensure that animals moved towards the same direction. Wherever it was necessary, corrugated plastic cardboard was positioned to increase animal contact with the obstacles and to prevent them from taking shortcuts (Fig 1D). The experimenter started each session by placing animals from both groups at the starting point. Once the mice reached the goal, they were allowed to consume the reward. Next, they were returned to the starting point and the reward was refilled. Prior to and after each training session, the entire track was wiped with alcohol.

Prior to the start of training, we familiarized the animal with the chocolate milk reward inside their homecage. In the first week, both groups were habituated on the empty track and learned to run on the track, which was gradually filled with ramps that they had to traverse to get to the goal location (reward). During the following weeks (from weeks two through nine), no alterations were made to the track for the control group, which continued to run over the ramps. In contrast, starting from the second week for the enriched group, the ramps were gradually replaced by three new obstacles every day until a total of 12 new obstacles filled the entire track. In the subsequent weeks, the enriched group experienced three track rearrangements across each session, which involved rotating or switching the positions of two obstacles every day, interleaved with two new obstacles. This paradigm continued until week nine, when the training was completed (Sup. Fig 1).

Over the course of each session, for the enrichment group, a new obstacle (such as seesaw, stairs, or tunnel) or track manipulation was added every ten minutes, consistently increasing the course’s level of difficulty. By the end of the last training week, the enriched animals had experienced 84 different obstacles. Additionally, every sixth training day, the enriched animals were exposed to a completely novel track (sets 1 to 7), while the control track remained unchanged throughout, to control for the effect of physical activity.

### Surgical Procedure

Following nine weeks of training, all animals received a cranial window implant over the dorsal cortex according to the same procedures as previously detailed in Esteves et al. (2021; 2023). Briefly, animals received buprenorphine (0.1 mg/kg, subcutaneous), dextrose with atropine (5 mL D5W; atropine 0.06 mg/kg, subcutaneous), and dexamethasone (2 mg/kg, intramuscular) to reduce cortical swelling. They were then anaesthetized with isoflurane (1% to 1.5%, O2: 1 l/min). The hair over the scalp was removed using a hair-clipper before placing the mice in a stereotaxic frame. Body temperature was regulated using a heating pad maintained at 37°C body temperature, regulated using a rectal probe attached to a temperature controller in a feedback loop. Animals’ eyes were protected with ophthalmic ointment. After placement in ear-bars, lidocaine (0.5%, 7 mg/kg) was injected subcutaneously under the incision site. The skin was disinfected with chlorhexidine-alcohol before the skull was exposed, and a custom-made titanium head-plate was fixed using adhesive cement (C&B-Metabond, Parkell). Using a high-speed dental drill with sterile drill bit, we performed a 5 mm diameter, circular craniotomy (Fig. 1F) over the dorsal cortex (AP: 2 mm anterior to −3 mm posterior from bregma; ML: -2.5 mm to +2.5 mm), with the dura left intact. Next, a window was implanted over the craniotomy, consisting of a 7 mm diameter round coverslip placed on top of two stacked 5 mm diameter coverslips. The 5 mm coverslips rested against the brain surface, while the edge of the 7 mm coverslip made contact with the animal’s skull. The three layers of coverslips were affixed to one another with optical adhesive (NOA71, Norland), and the window was attached over the craniotomy with tissue glue (Vetbond; 3M). The skin around the head-plate was sealed with tissue glue, and a layer of dental acrylic (Jet Tooth Shade Powder and Liquid, Lang Dental Manufacturing Co.) was applied between the window and the head-plate. At the end of the surgery, a rubber ring was affixed to the outer edge of the head-plate in order to create a well to hold water between the imaging region and the water immersion objective (Fig 1E). Postoperative analgesic treatment was continued with meloxicam (Metacam, 1 mg/kg, subcutaneous) and enrofloxacin (Baytril, 10 mg/kg, subcutaneous) for 3 days after surgery. All procedures were performed under aseptic conditions.

### Treadmill apparatus

After cranial window implantation, mice were given a week to recover from surgery before being water restricted. Mice under water restriction had free access to water in their home cages for 30 minutes each day following the recording sessions. Their weight was monitored to ensure that it remained above 85% of their baseline weight, which was established as the average weight over three days prior to the start of water restriction. Over the course of a week, animals were gradually accustomed to being head-fixed and to rest over the clamped treadmill. The treadmill consisted of a 150 cm long belt made from Velcro loops (Country Brook) with tactile cues placed at different locations (made from hot-glue, reflective tape, and Velcro). The belt was guided by two, 10-cm-diameter, polyamide wheels located at each end of the treadmill, and an optical encoder (Avago Tech) was mounted to the wheel shaft to track the movement of the animal. At the end of each lap, detected by a photoelectric sensor (Omron), a solenoid pinch valve (Bio-Chem) released a drop of sucrose water to the animal (10% sucrose water; 2.5µl in volume). Reward delivery was controlled by a custom-designed circuit and a microcontroller (Arduino Mega, Farnell).

### Two-photon microscopy and imaging pre-processing

Two-photon calcium imaging was conducted using a Thorlabs Bergamo II multi-photon microscope. GCaMP6s was excited at 920 nm (80–120 mW power at the objective) using a Ti:Sapphire femtosecond pulsed laser (Coherent Chamelion Ultra II). Emitted photons were gathered by a 16x water immersion objective (Nikon; NA = 0.8) and were detected with a GaAsP photomultiplier tube (Hamamatsu). The beam was raster-scanned by Galvo-Resonant scanners over a 835 μm × 835 μm square FOV, and the resulting frames were digitized to a resolution of 800 × 800 pixels at a rate of ∼19 Hz. Imaging data from all animals were consistently acquired from one hemisphere (either the left or right) at depths between 130µm to 190µm (layers II/III).

Image registration, cellular ROI detection, and activity extraction was performed using Suite-2P (Pachitariu et al. 2017), as previously described (Esteves et al. 2021; 2023). Cellular ROIs detected were then visually inspected and labelled as cells or non-cells by one experienced user, blind to the study group, based on morphological features and stereotypical fluorescence markers. The time-courses for each cell was obtained by first extracting the raw fluorescence traces (F) for each cellular ROI, calculated by subtracting the neuropil contamination from the mean fluorescence of all pixels within the ROI. Neuropil masks were estimated by separately dilating each ROI mask by 8 pixels and 1 pixel, and by subtracting the two to obtain an outer ring (Bonin et al. 2011). The raw neuropil fluorescence, computed as the mean fluorescence of the remaining ring mask, was then applied to reconstruct the neuropil fluorescence timeseries for each ROI. For each raw fluorescence signal (F), a rolling baseline (F0) was calculated using a Gaussian filter with a 10-second width, followed by minimum and maximum filtering over a 60-second window. Finally, the baseline-corrected fluorescence signal (ΔF/F0) was computed as (F−F0)/F0. Next, we inferred the spiking rate of each neuron by deconvolving the ΔF/F0 time-courses using constrained non-negative matrix factorization (Pnevmatikakis et al. 2016), and all subsequent analyses were conducted on the deconvolved time-courses using MATLAB (R2023a Mathworks, Natick, MA).

### Data analysis

The methods used for identifying spatially tuned neurons have been thoroughly documented in previous studies (Chang et al. 2020; Esteves et al. 2021). Briefly, in order to be successful classified as a spatially selective cell, a neuron must satisfy two criteria: First, the spatial information conveyed by a neuron about the animal’s location must be greater than the 95th percentile of a shuffled distribution (Skaggs, McNaughton, and Gothard 1992). Second, a continuous wavelet transform (Mexican Hat) was conducted over the spatial tuning curve of the neurons. If a local maximum exceeds 3 median absolute deviations from the wavelet coefficients at the lowest scale of the transform (where each scale corresponds to a spatial bin), the activity over the curve was identified as a potential place field (Chang et al. 2020; 2023). For each candidate place field, the width needs to be between 5% and 80% of the environment’s total length, the mean activity needs to be 2.5 times higher than the activity outside of the place field, and at least one-third of the individual trials must have their peak activity within the place field. If neurons exhibited at least one place field that met these requirements, they were classified as spatially selective.

Lifetime sparsity, which quantifies how dispersed a neuron’s firing during a recording session, was calculated as:

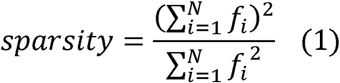

where *f*_i_ is the deconvolved ΔF/F_0_ activity in the i^th^ time frame over a total of N = 1000 frames (≈52 s). Population sparsity was quantified also using (1), however f_i_ is the deconvolved ΔF/F_0_ activity of the i^th^ neuron over a total of N cells. Sparsity ranges between 0 to 1, where neurons with sparser coding are represented by lower values, and neurons that are more active during the session are represented by higher values.

Explained Variance (EV) used to assess memory reactivation was calculated as:

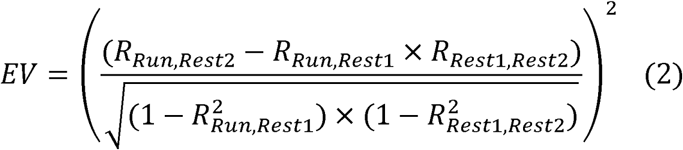

where the R-values correspond to the upper triangle of the Pearson correlation matrices for the deconvolved time-courses vectors of all neurons for each session. The Reverse Explained Variance (REV) was calculated by switching the position of Rest1 and Rest2 in the equation above (Hemant S. Kudrimoti, Barnes, and McNaughton 1999; Pennartz et al. 2004; Tatsuno, Lipa, and McNaughton 2006).

Only sessions in which the animals completed more than 10 laps were included in the results. For sessions with more than 20 laps, only the first 20 laps were used for data analysis. When possible, the imaging window was kept constant to image the same cells across days. To that end, we used blood vessels and specific cells as landmarks to align the FOV to that of the reference day at the beginning of each imaging session. To detect persistent cells across two imaging sessions, we first registered the binary masks of all detected cell ROIs between the two sessions by rigid transformation (rotation and translation). Then, neuron ROIs that showed more than 55% pixel overlap of the overall area (Jaccard distance) were identified as the same neuron. Recordings from days 4, 5, and 6 included the same ROIs across sessions and were used to assess neuronal stability over time. In contrast, recordings from days 1, 3, and 6 did not necessarily share the same ROIs across days, and analyses using these sessions focused on changes in cortical activity associated with task learning.

### Statistical analysis

All statistical tests performed in this work were conducted using MATLAB functions (R2017a Mathworks, Natick, MA) and IBM SPSS Statistics (version 28.0.1.0). Non parametric test were performed on datasets that didn’t pass the homogeneity of variance test (Levene’s test). Bonferroni correction was used to adjust for multiple comparisons when conducting post-hoc tests after a two-way ANOVA to compare all means from both groups. Further details of all statistical tests implemented in this study are provided in Table 1.

## Notes

### Competing Interest Statement

The authors have declared no competing interest.

